# Reward-driven enhancements in motor control are robust to TMS manipulation

**DOI:** 10.1101/2020.01.12.903419

**Authors:** Olivier Codol, Joseph M. Galea, Roya Jalali, Peter J. Holland

## Abstract

A wealth of evidence describes the strong positive impact that reward has on motor control at the behavioural level. However, surprisingly little is known regarding the neural mechanisms which underpin these effects, beyond a reliance on the dopaminergic system. In recent work, we developed a task that enabled the dissociation of the selection and execution components of an upper limb reaching movement. Our results demonstrated that both selection and execution are concommitently enhanced by immediate reward availability. Here, we investigate what the neural underpinnings of each component may be. To this end, we disrupted activity of the ventromedial prefrontal cortex and supplementary motor area using continuous theta-burst transcranial magnetic stimulation (cTBS) in a within-participant design (N=23). Both cortical areas are involved in reward processing and motor control, and we hypothesised that disruption of their activity would result in disruption of the reward-driven effects on action selection and execution, respectively. To increase statistical power, participants were pre-selected based on their sensitivity to reward in the reaching task. While reward did lead to enhanced perforance during the cTBS sessions and a control sham session, cTBS was ineffective in altering these effects. These results may provide evidence that other areas, such as the primary motor cortex or the premotor area, may drive the reward-based enhancements of motor performance.

## Introduction

In saccadic eye movements, reward has a well-known ability to invigorate motor control, enhance accuracy, and promote accurate action selection in the face of potential distractors (Kojima and Soetedjo, 2017; Manohar et al., 2015; Sohn and Lee, 2006; Takikawa et al., 2002). Recently, we extended these behavioural findings from eye movements to reaching movements (Codol et al., 2019). Specifically, we found that reward enhanced action selection by increasing participants’ propensity to move towards the correct target in the presence of a distractor target, while reaction times were not impeded. Execution of reaching movements also showed a pronounced increase in peak velocity (vigour) with reward, while radial accuracy was maintained. While these reward-driven improvements are now behaviourally well-characterised and confirmed in a number of previous reports (Griffiths and Beierholm, 2017; Reppert et al., 2018; Summerside et al., 2018), the neural substrates of these effects remain unknown.

During a sensorimotor task, a stream of information contributes to the generation of movement, travelling from visual and proprio-tactile sensory afferents to high-level prefrontal and parietal associative areas; then forming into a motor plan in the supplementary motor area (SMA) and pre-motor cortices, to finally produce a motor command which travels from the primary motor cortex (M1) to the spinal cord and to the effector muscles (Castiello, 2005; Hikosaka et al., 2002; Shadmehr and Krakauer, 2008; Thorpe and Fabre-Thorpe, 2001). Therefore, to pin down the neural substrates of reward-driven enhancements, one can ask at which point of this sensory-prefrontal-premotor-motor loop does reward influence the processing stream. To this end, we aimed to disrupt the activity of specific cortical regions in the sensorimotor pathway to establish a causal relationship with behaviour. We applied continuous theta-burst transcranial magnetic stimulation (cTBS; Huang et al., 2005; Zenon et al., 2015) immediately prior to participants performing the task. Such manipulation has been shown to disrupt neural activity for 20 min following cessation of stimulation (Huang et al., 2005). The targeted regions would therefore continue to be disrupted for the entire duration of our behavioural task, without the need to stimulate during task performance.

Some evidence to determine potential cTBS targets may come from the literature on attentional processes, as sensitivity of attention to reward is a well-known phenomenon (Sarter et al., 2006). For instance, reward-driven selection improvements similar to our observations have been reported in the Eriksen flanker task (Hübner and Schlösser, 2010), a seminal paradigm for studying attentional capacity. The authors argued that reward may trigger an enhancement of sensory information encoding, drastically improving evidence accumulation and thus action selection. Physiologically, such a mechanism has been shown in rats to occur in visual cortices through cholinergic modulation (Goard and Dan, 2009; Pinto et al., 2013), and imaging studies show that occipital regions exhibit the most sensitivity to reward in attentional tasks in humans (Anderson, 2016; Tosoni et al., 2013). Thus, it may be that the reward-driven selection improvements we report are due to early enhancement of visual sensory processing in the sensorimotor loop. This possibility has also been raised in a study of saccades (Manohar et al., 2015). However, in that study, the authors also found that Parkinson’s disease patients did not exhibit the increase in selection accuracy with reward seen in healthy aged-matched controls.

These results suggest that though acetylcholine may play a role in enhancement of selection accuracy, a role for dopamine should be considered as well. In line with this argument, a large number of imaging studies have demonstrated the involvement of the posterior and anterior cingulate cortices, and ventro-medial prefrontal cortex (vmPFC) in reward processing (Blair et al., 2013; Daw et al., 2005, 2006; Graybiel, 2008; Klein-Flugge et al., 2016), regions that are heavily dependent on dopamine innervation (Arnsten, 1998) and also involved in the sensorimotor loop (Hikosaka et al., 2002). Furthermore, imaging evidence shows that vmPFC encodes the value of different stimuli during a decision-making task involving motor effort (Klein-Flugge et al., 2016). Consequently, prefrontal or occipital areas could both be considered as potential targets for cTBS. However, since occipital areas are not only involved in reward processing but also a large array of core visual functions, cTBS in these regions could potentially disrupt basic motor performance, and thus expose any results to unnecessary confounds. Therefore, we focus on prefrontal cTBS manipulations in this study to assess action selection susceptibility to reward. While the anterior cingulate cortex and the vmPFC are both possible candidates, the anterior cingulate cortex cannot be stimulated using cTBS due to its deep location, we therefore tested our hypothesis by targeting the vmPFC.

Regarding execution, reward-based improvements may be similarly due to enhanced encoding of visual information, thereby allowing more vigorous movements at no accuracy cost. However, this would not explain the reward-driven increase in feedback control (Carroll et al., 2019; Manohar et al., 2019) and end-point stiffness we observe (Codol et al., 2019). Rather, reward could directly modulate M1, as M1 activity has been shown to be highly sensitive to reward (Bundt et al., 2016; Galaro et al., 2019; Kapogiannis et al., 2008; Mawase et al., 2016, 2017; Ramkumar et al., 2016; Thabit et al., 2011; Zhao et al., 2018), shaping processing near the end of the sensorimotor arc. Another reasonable hypothesis is that reward information is integrated earlier on, with M1 being merely the final recipient. Several prefrontal regions upstream of M1 are involved in action planning, including the SMA, a region also showing strong sensitivity to reward (Klein-Flugge et al., 2016; Stanford et al., 2013; Zenon et al., 2015). In Parkinson’s disease patients, who express apathy symptoms sometimes interpreted as a lack of vigour, also show altered SMA activity (Hendrix et al., 2018; Rascol et al., 1994). In recent work, it was argued that SMA encodes sensitivity to effort (Klein-Flugge et al., 2016), which is hypothesised to drive the change in vigour during motor control (Manohar et al., 2015; Mazzoni et al., 2007). While cTBS stimulation over M1 would not answer whether reward is integrated in M1 or earlier on, SMA stimulation could provide more conclusive evidence. If an effect on reward-driven enhancement of execution performance is seen, this would confirm that reward information is indeed integrated earlier than might be initially expected for reaching movements (Mawase et al., 2016, 2017; Thabit et al., 2011).

Consequently, the aim of this study was first to replicate previously reported findings regarding the effect of reward on this reaching task; and second, to alter the effect of reward on action selection and action execution through cTBS of the vmPFC and SMA, respectively.

## Methods

### Participants

26 of 34 screened participants (see “screening session” section for details) were selected based on their performance on the reaching task. Of those 26 selected participants, one was excluded due to medical reasons, and two participants retracted after the second session. Therefore, 23 participants (median age: 22, range: 18-39, 15 female) took part in the experiment and were remunerated £15/hour in addition to performance-based monetary rewards during the reaching task. All participants were right-handed, free of epilepsy, familial history of epilepsy, motor, psychological or neurological conditions, or any medical condition forbidding the use of cTBS or MRI. The study was approved by and completed in accordance with the University of Birmingham Ethics Committee.

### Task design

The behavioural task was identical to the first experiment of Codol et al. (2019), except that only 0p (pence) and 50p trials were used. Participants performed the tasks on an end-point KINARM (BKIN Technologies, Ontario, Canada). They held a robotic handle that could move freely on a horizontal plane in front of them, with the handle and their hand hidden by a panel (figure 1A). The panel included a mirror that reflected a screen above it, and participants performed the task by looking at the reflection of the screen (60Hz refresh rate), which appeared at the level of the hidden hand. Kinematics data were sampled at 1kHz.

**Figure 1:**
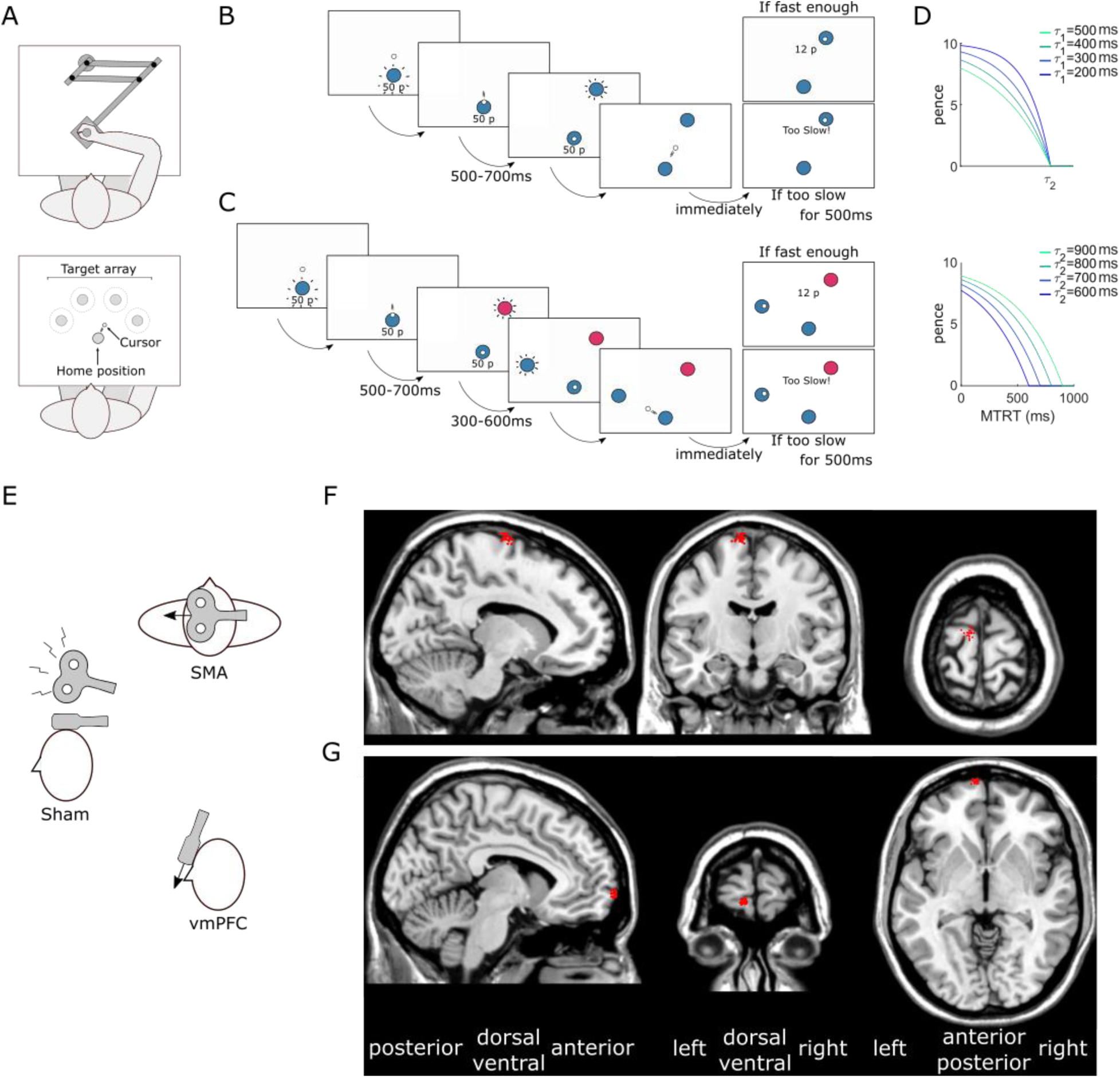
Behavioural experiment and cTBS procedure. A. Participants reached to a series of targets using a robotic manipulandum. B. Time-course of a normal trial. Participants reached at a single target and earned money based on their performance speed (sum of movement time and reaction time; MTRT). If they were too slow (MTRT<*τ*_2_), a message “*Too slow!*” appeared instead of the reward information. Transition times are indicated below for each screen. A uniform distribution was employed for the transition time jitter. C. Time-course of a distractor trial. Occasionally, a distractor target appeared, indicated by a colour different from the starting position. Participants were told to wait for the second, correct target to appear and reach toward the latter. D. The reward function (here for a 10p trial) varied based on two parameters *τ*_1_ (upper plot; *τ*_2_ fixed at 800ms) and *τ*_2_ (lower plot; *τ*_1_ fixed at 400ms). E. position of the cTBS coil(s) relative to the head in each of the 3 conditions. The black arrows represent the current orientation. F. Sagittal, coronal and axial planes of an MNI-normalised brain scan (*ch2.nii.gz* in MRIcron). The red dots indicate each participant’s SMA stimulation sites. G. vmPFC stimulation sites.

Each trial started with the robot handle bringing participants to a point 4cm in front of a fixed starting position. A 2cm diameter starting position (angular size ∼3.15°) then appeared, with its colour indicating the reward value of that trial. The reward value was also displayed in 2cm-high text (angular size ∼3.19°) under the starting position (figure 1B-C). Because colour luminance can affect salience and therefore detectability, luminance-adjusted colours were employed (see http://www.hsluv.org/) and colours assigned to distractors or real targets were counterbalanced across participants. For a given participant, the two colours coding for the real targets were never the same as the two colours coding for distractor targets.

From 500 to 700ms after participants entered the starting position (on average 587±354ms after the starting position appeared), a 2cm diameter target (angular size ∼2.48°) appeared 20cm away from the starting position, in the same colour as the starting position. Participants were instructed to move as fast as they could towards it and stop in it. They were informed that a combination of their reaction time and movement time defined how much money they would receive, and that this amount accumulated across the experiment. They were also informed that end-position was not factored in as long as terminated the movement within 4cm of the target centre. There were 4 possible target locations positioned every 45° around the midline of the workspace, resulting in a 135° span (figure 1A).

The reward function was of a closed-loop design that incorporated the recent history of performance, to ensure that participants received similar amounts of reward despite idiosyncrasies in individuals’ reaction times and movement speed. Furthermore, the closed-loop nature of the reward function ensured that the task remained consistently challenging over the course of the experiment (Berret et al., 2018; Manohar et al., 2015; Reppert et al., 2018). To that end, the reward function was defined as:

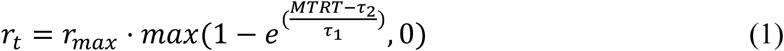

where *r*_*max*_ was the maximum reward value for a given trial, *MTRT* the sum of reaction time and movement time, and *τ*_1_ and *τ*_2_ adaptable parameters varying as a function of performance (figure 1D). Specifically, *τ*_1_ and *τ*_2_ were the median of the last 20 trials’ 3-4th and 16-17th fastest MTRTs, respectively, and were initialised as 400 and 800ms at the start of each participant training block. *τ* values were constrained so that *τ*_1_ < *τ*_2_ < 900 was always true. In practice, all reward values were rounded up to the nearest penny so that only integer penny values would be displayed.

Targets were always of the same colour as the starting position (figure 1B), but occasional distractor targets appeared, indicated by a different colour than the starting position (figure 1C). Participants were informed to ignore these targets and wait for the second target to appear. Failure to comply resulted in no monetary gain for this trial. The first target (distractor or not) appeared 500-700ms after entering the starting position using a uniform random distribution, and correct targets in distractor trials appeared 300-600ms after the distractor target using the same distribution.

When reaching movement velocity passed below a 0.3 m/s threshold, the end position was recorded, and monetary gains were indicated at the centre of the workspace. After 500ms, the robotic arm then brought the participant’s hand back to the initial position 4cm above the starting position.

### Procedure

The experiment took place over five sessions, with a gap of at least five days between sessions. The first session was a screening session, in which participants were selected based on their performance during the behavioural task. In the second session, a structural MRI scan of each participant’s brain was acquired, and used for the third to fifth session, during which participants performed the behavioural task after receiving either sham, SMA or vmPFC cTBS (figure 1A). The order of stimulation was pseudo-randomly counterbalanced across participants. Before every session, participant’s health condition was assessed in accordance to the guidelines of the Ethics Committee of the University of Birmingham (UK).

### Screening session

In the first session, participants were first screened for medical or psychological conditions that could exclude them from the study. They were then introduced to the cTBS technique by reading a leaflet, and they could ask any questions they wished to the experimenter. Next, they were exposed to theta-burst stimulation on their forearm to get acquainted with the sensation of stimulation. Their active motor threshold (AMT) was then determined by finding the minimal single-pulse transcranial magnetic stimulation intensity on M1 that resulted in the visible contraction of the first dorsal interossei (FDI) muscle of the preactivated right hand in 5 out of 10 trials. Finally, participants performed the behavioural task.

Participants first practiced the task in a 48 trials training block with a 25p trial value. They were informed that money obtained during the training would not count toward the final amount they would receive. The starting position and target colours were all grey during training. They then performed a 16 trials, distractor-free baseline block with 0p and 50p trials and were informed that their score now counted toward their final monetary gain. Finally, they experienced a 224 trials main block that included 96 (42.9%) distractor-containing trials randomly interspaced. For the three cTBS sessions, participants repeated the same task, with the exception of the training block which was removed. Because this study aimed to manipulate a previously characterised effect, participants were selected for the subsequent sessions only if they showed a reward driven increase in both peak velocity and selection accuracy.

Using the resulting behavioural data, participants were then screened for an effect of reward on execution and selection accuracy. Specifically, participants were expected to show an increase in peak velocity and selection accuracy (*i.e.* increased propensity to ignore a distractor target) in rewarded trials compared to non-rewarded trials. Participants who did not show both of these effects or showed an overly weak effect were excluded. Of note, no participant showed an effect opposite to the effect of interest.

### cTBS procedure

Using a 3-T Philips (Amsterdam, The Netherlands) scanner, high-resolution T1-weighted images were acquired for each participant (1×1×1mm voxel size, 175 slices in sagittal orientation). The image was then normalised to an MNI template using an affine (12 parameter) transformation (Jenkinson and Smith, 2001; Jenkinson et al., 2002) with the software Statistical Parametric Mapping 12 (SPM12, London, UK). Regions of interest were then marked using MRIcron (Rorden and Brett, 2000). The MNI coordinates used were *x* = −8/*y* = −9/*z* = 77 for the SMA and −7/71/−4 for the vmPFC (figure 1B, C). More specifically, the SMA target region was the posterior part of the superior frontal gyrus, or the most prominent posterior part of Brodmann area 6 (Arai et al., 2012; Zenon et al., 2015); the vmPFC target region was the most anterior part of medial orbitofrontal gyrus, or Brodmann area 10 near the limit with Brodmann 11 (Blair et al., 2013; Lev-Ran et al., 2012). These positions were all in the left hemisphere (Arai et al., 2012; Lev-Ran et al., 2012) since all our participants were right-handed. The marked scans were then transformed back into their original space using each participant’s inverse transform with SPM12, and the position of each mark was manually inspected and adjusted to the closest location minimising distance between the target position and the scalp (Galea et al., 2010; Huang et al., 2005), giving subject-specific target locations. The resulting marked individual scans were then imported to a BrainSight 2 neuronavigation system (Rogue Research Inc, Montreal, Quebec), and each region of interest was targetted with the TMS coil using its motion-capture tracking function. The SMA stimulation was performed at −90° from the midline and the vmPFC stimulation was performed at 0° from the midline, with the coil being placed tangencially to the forefront (*i.e.* almost vertically for the vmPFC, see figure 1E).

cTBS was applied with a figure-of-eight, 80mm diameter coil (Magstim Co Ltd, Whitland, UK). We employed the continuous theta-burst stimulation technique, with one cycle lasting 40s, at 80% AMT or 48% intensity, whichever was the lowest. A total of 200 burst trains were applied at a frequency of 5Hz, with 3 pulses per burst and a pulse frequency of 50Hz—giving a total amount of 600 pulses. These parameters were all based on Huang et al. (2005) and Galea et al. (2010). During all cTBS sessions (including the sham session), participants were asked if they felt fine immediately after the stimulation was performed, and upon confirmation, were asked to move approximately two meters from the stimulation chair to the chair on which they could perform the behavioural task.

### Data analysis

The pre-registered *a priori* hypotheses, cTBS procedure, dataset and analysis scripts are all available online on the Open Science Framework website (https://osf.io/tnkrj/). Analyses were performed using custom Matlab scripts (Matworks, Natick, MA). Bayesian analyses were performed using JASP (JASP, Amsterdam, The Netherlands).

Trials were manually classified as distracted or non-distracted. Trials that did not include a distractor target—*i.e.* no-distractor trials—were all considered non-distracted. Distracted trials were defined as trials where a distractor target was displayed, and participants initiated their movement (*i.e.* exited the starting position) toward the distractor instead of the correct target. If participants readjusted their reach “mid-flight” to the correct target or initiated their movement to the correct target and readjusted their reach to the distractor, this was still considered a distracted trial.

Reaction times were measured as the time between the correct target onset and when the participant’s distance from the centre of the starting position exceeded 2cm. In trials that were marked as “distracted” (*i.e.* participant initially went to the distractor target), the distractor target onset was used. In distractor-containing trials, the second, correct target did not require any selection process to be made, since the appearance of the distractor target informed participants that the next target would be the correct one. For this reason, reaction times were biased toward a faster range in non-distracted trials. Consequently, mean reaction times were obtained by including only no-distractor trials, and distracted trials. For every other summary variable, we included all trials that were not distracted trials, that is, we included non-distracted trials and no-distractor trials.

Trials with reaction times higher than 1000ms or less than 200ms, and non-distracted trials with radial errors higher than 6cm or angular errors higher than 20° were removed. Overall, this accounted for 0.49% of all trials. Speed-accuracy functions were obtained for each participant by binning data in the *x*-dimension into 50 quantiles and averaging all *y*-dimension values in a *x*-dimension sliding window of a 30-centile width (Manohar et al., 2015). Then, each individual speed-accuracy function was averaged by quantile across participants in both the *x* and *y* dimension.

### Statistical analysis

In the pre-registration of this study, we indicated that group statistics would be performed using a 2×3 repeated-measure ANOVA, with reward value (0p versus 50p) as the first factor, and cTBS group (sham, SMA, vmPFC) as the second factor. However, because main effects were only detected in the first factor (0p-50p) and no effect was found in the cTBS condition, we also performed *post-hoc* Bayesian analyses to assess the evidence in favour of the null hypothesis regarding cTBS manipulation. Results were identical regarding significant effects in the frequentist versus Bayesian approach. Frequentist ANOVAs were performed in MatLab, and Bayesian statistics were done using the Bayesian repeated-measure ANOVA function in JASP with mean summary statistics pre-computed and exported as csv files using MatLab. Results are reported as Bayes factors for each model against the null model (BF_10_) and for each model against the best model (BF_best_). A BF of 1 indicates that there is no evidence in favour of the null or the alternative model, *i.e.* the data is ambiguous (Wagenmakers et al., 2011). A BF that thends toward 0 indicates increasing evidence toward the null model, and inversely, a BF that tends toward +∞ indicate stronger evidence for the alternative. Note that this is a log-scale, *i.e.* a BF of 2 is as much evidence for the alternative model than a BF of 0.5 is for the null (Wagenmakers et al., 2011).

The default prior parameters were used, i.e. a Cauchy prior with r-scale of 0.5 for fixed effects (there was no random effect or covariate). Sampling values for numerical accuracy and model-averaged posteriors were left in the “automatic” position. To obtain model-averaged posteriors, the posterior density function of a given factor level coefficient must be averaged across all models, with each posterior density being weighted by the probability of its respective model. In other words, it is a weighted mean of posterior effects across all models. For all plotted variables, bootstrapped 95% confidence intervals of the mean were obtained using 10,000 permutations.

## Results

Similar to Codol et al. (2019), reward improved both the selection and execution components of reaching movements (figure 1). Specifically, reward led to faster reaction times (*F*(1,22) = 8.18, *p* = 0.009, partial *η*^2^ = 0.37; figure 1A), whilst also improving selection accuracy (*F*(1,22) = 16.7, *p* < 0.001, partial *η*^2^ = 0.76; figure 1B), clearly demonstrating that the selection component benefited from the presence of reward. Of note, the decrease of reaction times with reward in this study is surprising, as no significant effect had been observed in the same task in a previous study – though a non-significant trend in that direction could be observed (Codol et al. 2019). Regarding execution, peak velocity increased with reward (*F*(1,22) = 42.4, *p* < 0.001, partial *η*^2^ = 1.93; figure 1C) whilst movement time decreased (*F*(1,22) = 24.0, *p* < 0.001, partial *η*^2^ = 1.09; figure 1D). In addition, radial error (*F*(1,22) = 2.88, *p* = 0.10, partial *η*^2^ = 0.13; figure 1E) and angular error (*F*(1,22) = 2.98, *p* = 0.10, partial *η*^2^ = 0.14; figure 1F) were similar across rewarded and non-rewarded trials.

In contrast, while we expected to observe an effect of cTBS on the reward-driven effects, we observed no main effect or interaction effects for cTBS: reaction times (cTBS: *F*(2,44) = 0.05, *p* = 0.95, partial *η*^2^ = 0.002; interaction: *F*(2,44) = 0.65, *p* = 0.53, partial *η*^2^ = 0.03; figure 1A), selection accuracy (main effect of cTBS: *F*(2,44) = 0.40, *p* = 0.70, partial *η*^2^ = 0.02; interaction: *F*(2,44) = 1.12, *p* = 0.33, partial *η*^2^ = 0.05; figure 1B), peak velocity (cTBS: *F*(2,44) = 0.85, *p* = 0.43, partial *η*^2^ = 0.04; interaction: *F*(2,44) = 0.19, *p* = 0.83, partial *η*^2^ = 0.008; figure 1C), movement times (cTBS: *F*(2,44) = 0.21, *p* = 0.81, partial *η*^2^ = 0.009; interaction: *F*(2,44) = 0.78, *p* = 0.46, partial *η*^2^ = 0.03; figure 1D), radial (cTBS: *F*(2,44) = 0.79, *p* = 0.46, partial *η*^2^ = 0.04; interaction: *F*(2,44) = 1.08, *p* = 0.35, partial *η*^2^ = 0.05; figure 1E) and angular error (main effect of cTBS: *F*(2,44) = 1.18, *p* = 0.32, partial *η*^2^ = 0.05; interaction: *F*(2,44) = 0.16, *p* = 0.86, partial *η*^2^ = 0.007; figure 1F). This suggests that cTBS over vmPFC or SMA had no effect on behaviour. However, since the frequentist approach has inherent limitations regarding evidence for or against null effects, we performed *post-hoc* Bayesian analyses on our behavioural variables, and results are reported in table 1 for each model considered.

**Table 1:** Bayesian model comparison for kinematics variables. All models include participants as a random variables. The propability *p(M|data)* of a model given our dataset indicates which model is most likely compared to all other models considered. The most likely model for each variable is highlighted in bold. Bayesian factors (BF_10_) are the ratio between posterior likelihood of the model and the null (empty) model. A BF_10_ > 1 indicates that the model is more likely than the alternative null model. The BF_best_ row indicates the Bayes factor with respect to the best model.

| Model | Y ~ | Null (incl. pt.) | Reward | cTBS | cTBS + Reward | cTBS * Reward |
| --- | --- | --- | --- | --- | --- | --- |
| reaction times | $p(M/data)$ | 0.051 | **0.871** | 0.004 | 0.064 | 0.009 |
| | $BF_{10}$ | 1 | 16.966 | 0.076 | 1.253 | 0.180 |
| | $BF_{best}$ | 0.059 | 1 | 0.004 | 0.074 | 0.011 |
| selection accuracy | $p(M/data)$ | 1.150e -5 | **0.893** | 1.046e -6 | 0.089 | 0.018 |
| | $BF_{10}$ | 1 | 77668.98 | 0.091 | 7734.165 | 1562.092 |
| | $BF_{best}$ | 1.288e -5 | 1 | 1.171e -6 | 0.1 | 0.02 |
| peak velocity | $p(M/data)$ | 9.977e -9 | **0.786** | 1.759e -9 | 0.191 | 0.023 |
| | $BF_{10}$ | 1 | 7.881e +7 | 0.176 | 1.912e +7 | 2.299e +6 |
| | $BF_{best}$ | 1.269e -8 | 1 | 2.238e -9 | 0.243 | 0.029 |
| movement times | $p(M/data)$ | 1.168e -6 | **0.906** | 1.029e -7 | 0.082 | 0.012 |
| | $BF_{10}$ | 1 | 775804 | 0.088 | 70418.68 | 9941.661 |
| | $BF_{best}$ | 1.289e -6 | 1 | 1.136e -7 | 0.091 | 0.013 |
| radial error | $p(M/data)$ | **0.532** | 0.285 | 0.112 | 0.06 | 0.011 |
| | $BF_{10}$ | 1 | 0.535 | 0.21 | 0.113 | 0.022 |
| | $BF_{best}$ | 1 | 0.535 | 0.21 | 0.113 | 0.022 |
| angular error | $p(M/data)$ | **0.473** | 0.302 | 0.124 | 0.089 | 0.012 |
| | $BF_{10}$ | 1 | 0.639 | 0.262 | 0.188 | 0.024 |
| | $BF_{best}$ | 1 | 0.639 | 0.262 | 0.188 | 0.024 |

Comparing the candidate models using BF_10_, we see that the evidence in favour of the reward-only model is highest for both reaction times (BF_10_=16.9) and selection accuracy (BF_10_=7.76e+4), as well as peak velocity and movement time (BF_10_=7.88e+7 and 7.75e+5). These results are in line with the earlier frequentist analyses. In contrast, while the evidence pointed toward the null model for radial and angular error, the evidence against the reward-only model was weak, with BF_10_=0.53 and BF_10_=0.63, respectively. According to Wagenmakers et al. (2011), this represents only “anecdoctal” evidence for the null, emphasising the inconclusiveless of this result. Specifically, angular and radial accuracy were slightly lower in the reward condition compared to no reward (figure 1E-F).

To assess the impact of cTBS on performance, we included BF_10_ for three additional candidate models: y∼cTBS, y∼cTBS+reward, and y∼cTBS*reward (including an interaction). However, as a natural consequence of the strong evidence in favour of a reward effect, the BF_10_ of all variables tended to be very low for the cTBS-only model and extremely high for the models that included reward. To account for this, we compared Bayes factors with respect to the best model rather than the null model (BF_best_), which is tantamount to assessing how close the evidence for the considered model and best model is. Note that this method is uninformative for radial and angular error, because the null model is already the best, and because the BF_10_ remains weak (*i.e.* anecdotal) for all models anyway.

The cTBS+reward model exhibited strong evidence toward the null for all variables, except peak velocity, for which evidence toward the null was still strong (BF_best_=0.176) but less compelling than for the other variables. To assess which cTBS condition may drive this lower BF_best_, we assessed the model-averaged posterior distribution of each condition’s β coefficient (figure 3). Posterior effect sizes with respect to cTBS (figure 3A) indicate that this may be due to a small deviation of the SMA group effect size compared to sham and vmPFC. In comparison, the posterior effect size for reward showed a strong contrast between 0p and 50p (figure 3B), as expected from the high BF_10_ for the reward-only model (table 1). To assess whether there was an indirect impact of cTBS on reward-driven effects, we also considered the full cTBS*reward model. However, there was consistent and extreme evidence against this model compared to the best model for all variables considered (all BF_best_<0.03), excluding the possibility that cTBS manipulation had an impact in this task, directly or on the reward-driven effect. Illustrating this strong evidence against a potential interaction on peak velocity, the posterior coefficients for interactions were entirely overlapping (figure 3C).

**Figure 2:**
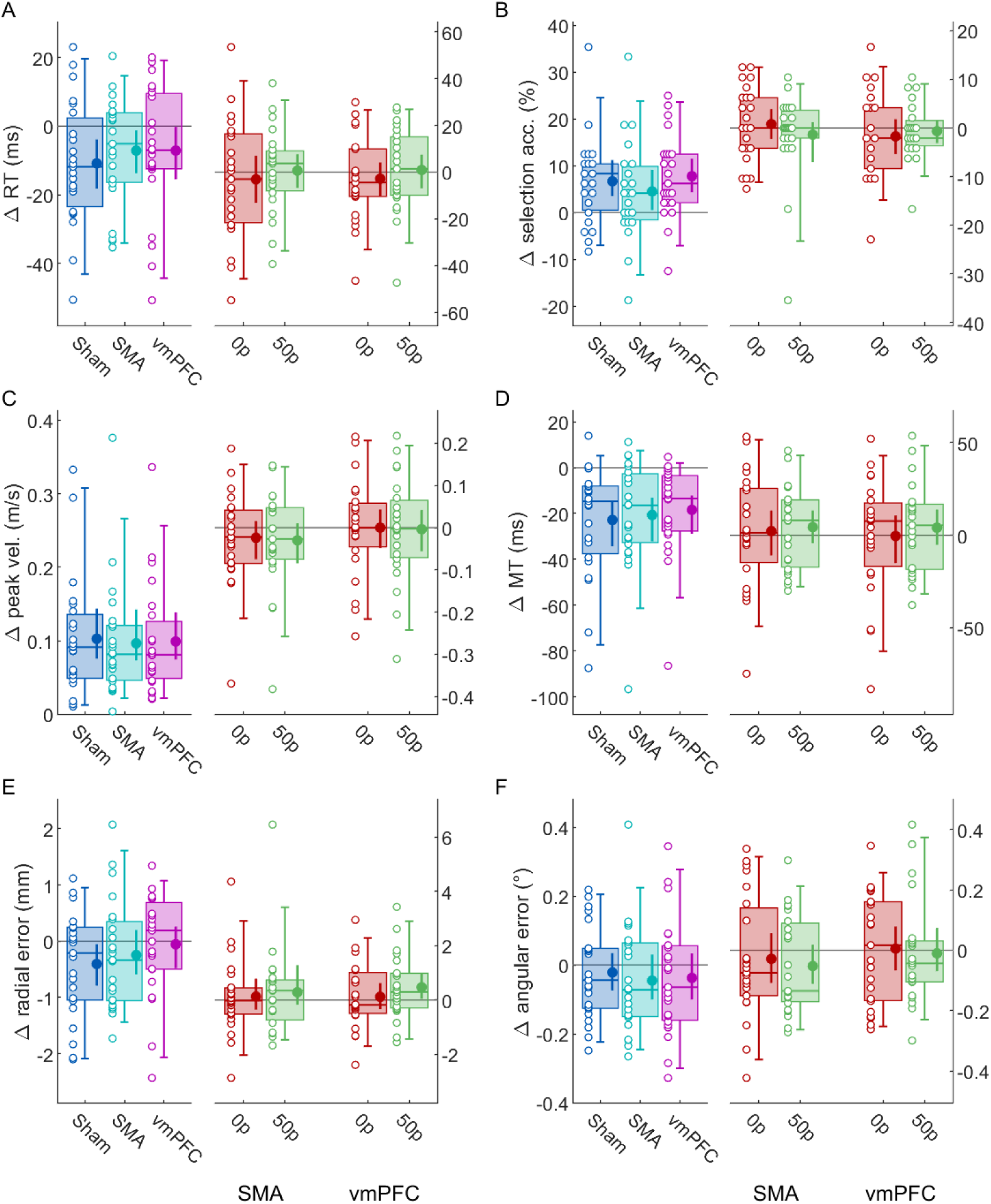
Effect of reward and cTBS on different behavioural variables. A. Reaction times. On the left, 50p trials performance for each cTBS group are normalised to 0p trials (*i.e.* reward-normalised), and on the right 0p and 50p trials for each cTBS group are normalised to sham performance (*i.e.* sham-normalised). The empty dots represent individual values for each group and the box plots indicate the [5-25-50-75-95] percentiles. The filled dot and and the error bars indicate the mean and bootstrapped 95% CIs of the mean. B-F. Other variables in the same format as panel A.

**Figure 3:**
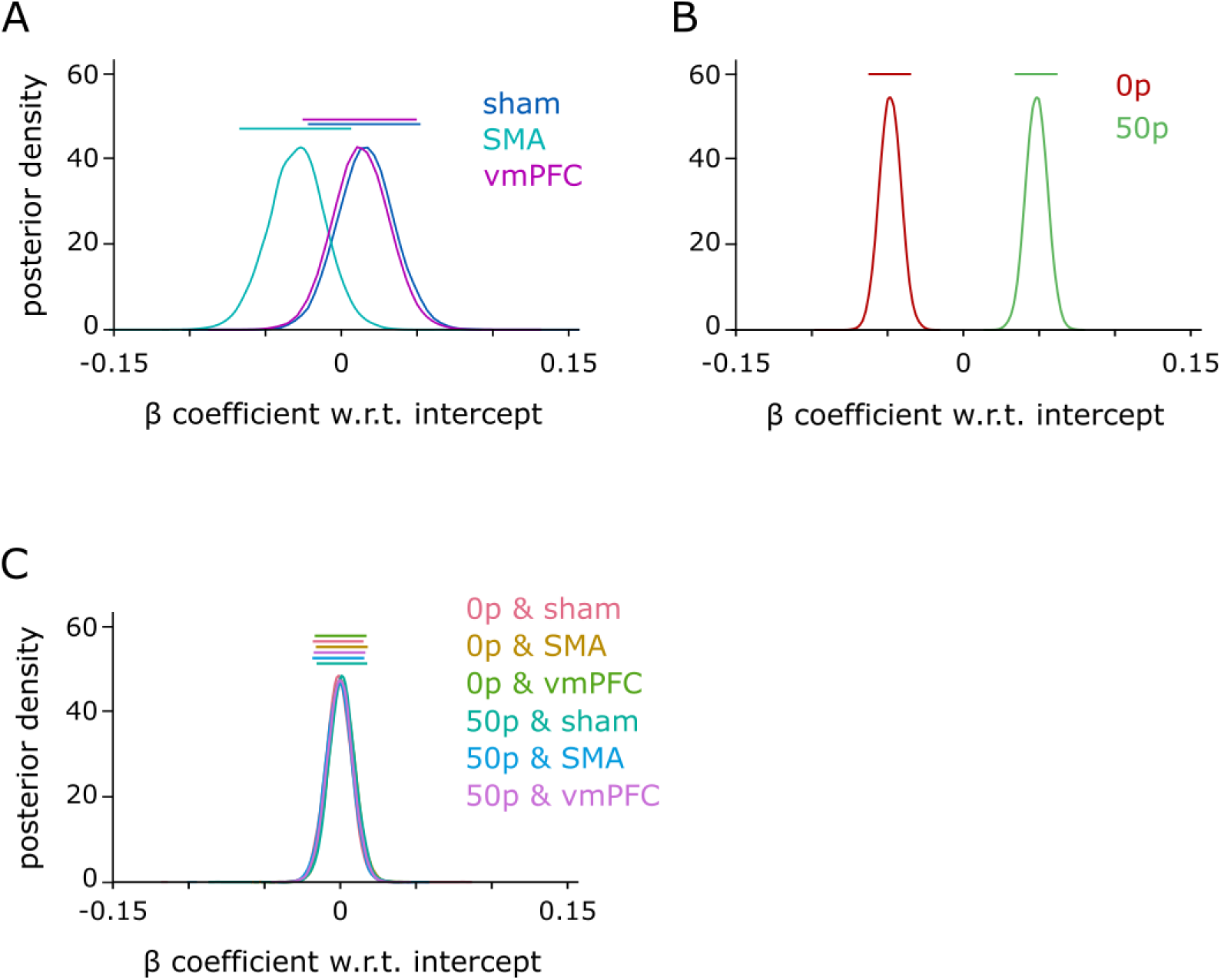
model-averaged posterior β coefficients for peak velocity. All effects are centred on 0 because they are experessed as a function of the model’s intercept. Bars on top of each probability density function indicate the 90% highest density interval. A. Posterior distributions for each cTBS condition. B. for each reward condition. C. for each possible interaction.

### Effect of reward and cTBS on speed-accuracy functions

Next, we assessed the speed-accuracy functions of the selection and execution components in all cTBS conditions. As can be seen in figure 4, we can consistently see a shift in the speed-accuracy functions of both these components with reward, in line with previous results (figure 4A-F). However, the execution speed-accuracy function in the SMA cTBS group does not exhibit a normal profile at baseline (0p trials; figure 4E). Instead, radial error appears to be maintained across the range of peak velocities displayed. However, this profile did not extend to rewarded trials. Because this behaviour at baseline is surprising, we examined individual speed-accuracy profiles for this condition to ensure this was not driven by outliers. We can observe from figure 5 that indeed, two participants displayed more accurate performance at high speeds for 0p trials in the SMA cTBS condition (middle panel), compared to the majority of participants. However, overall, there were also more participants who exhibited more accurate performance at higher speeds in this condition than in comparable conditions, such as 0p trials in the sham condition (figure 5, left panel) or the 50p trials in the SMA cTBS condition (right panel). Therefore, while no clear speed-accuracy trade-off was observed for the 0p trials in the SMA cTBS condition, it cannot be conclusively stated that this was driven by outliers. A possible reason for this unexpected result is that it is driven by the small, noisy trend observed for peak velocities illustrated in figure 3A. However, as demonstrated by the Bayes factor for peak velocity, this result remains too marginal to draw any strong conclusion.

**Figure 4:**
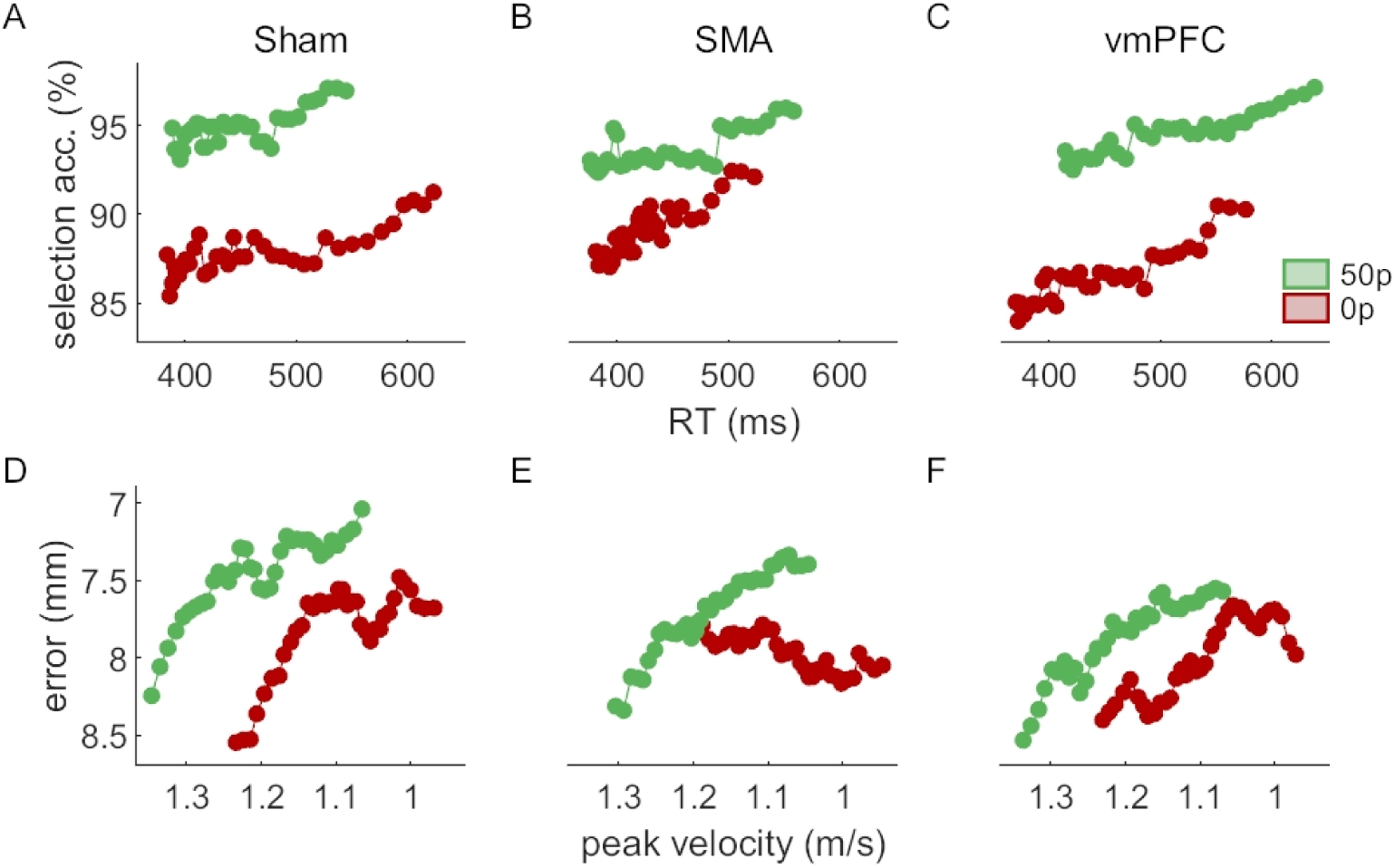
Speed-accuracy functions for each reward and cTBS condition. The selection (A-C) and execution (D-F) speed-accuracy functions are the top three and bottom three panels, respectively. The functions are obtained by sliding a 30% centile-wide window over 50 quantile-based bins and averaging each bin across participant. For the selection panels, the count of non-distracted trials and distracted trials for each bin was obtained, and the ratio (100*non-distracted/total) calculated afterwards. Note that the axes of the execution functions are reversed so that high speed and low accuracy are on the bottom-left corner like for the selection functions.

**Figure 5:**
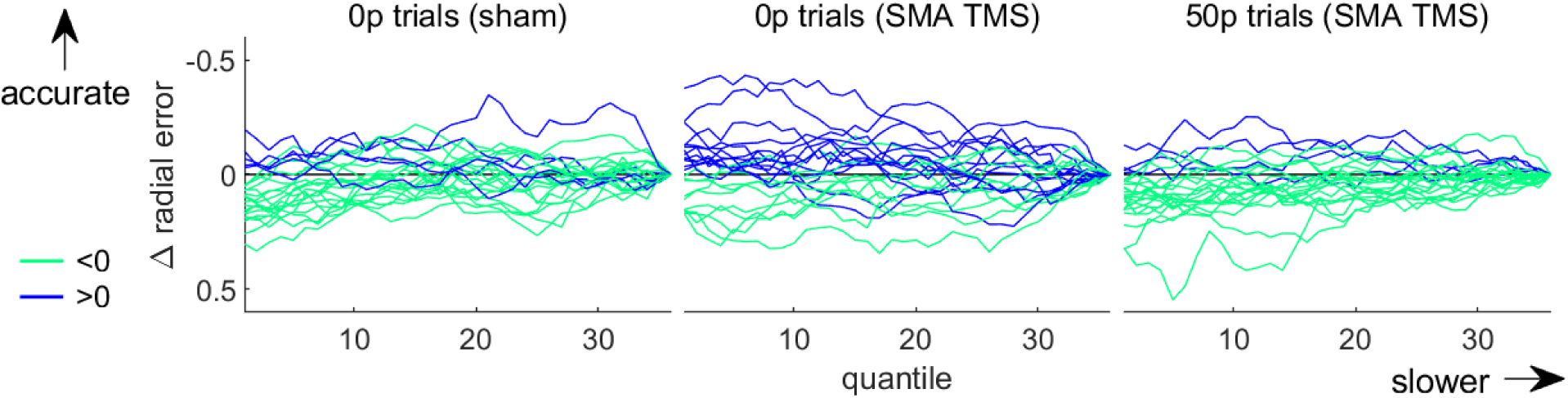
Individual speed-accuracy functions for the no-reward condition of the SMA cTBS group (middle) and for two control groups (right and left). The functions are obtained by sliding a 30% centile window over 50 quantile-based bins. Each individual profile is normalised to its end value. Profiles exhibiting an increase and a decrease in accuracy with slower movements are plotted in light green and blue, respectively.

## Discussion

In this study, we employed cTBS with the aim of perturbing activity in the vmPFC and SMA in order to modulate previously characterised reward-driven effects on selection and execution performance in a reaching task. While the effects of reward characterised in Codol et al. (2019) were reliably reproduced within participants and across a series of four sessions held on different days, cTBS stimulation of either of the two target regions did not result in any alteration of these effects.

The replication of reward-driven effects on a reaching task across weekly sessions and on the same individuals confirms the conclusions from our previous study (Codol et al., 2019). While it could be argued that this is natural considering that we pre-selected participants, it was not granted that an effect found on one day for a given participant could replicate consistently in a subsequent session held on another day. Nevertheless, one divergent result is that in this study we observe a reduction in reaction times with reward, in Codol et al. (2019) no significant effect had been observed despite a larger sample size (N=30). However, a similar trend that failed to reach significance had been observed. Here, pre-selecting participants may have allowed that trend to reach a significance threshold, suggesting that there is an effect of reward on reaction times, although it is likely a small effect size.

Interpreting the absence of any cTBS impact of the reward-driven effects is less straightforward, as drawing conclusions on the sole basis of non-significant results is a well-established fallacy (Altman and Bland, 1995). To gain a better understanding of the data, we performed a series of *a posteriori* Bayesian ANOVA analyses, allowing us to determine if the non-significant results are actually null results. However, this does not negate the inconclusive nature of a null result *per se*. Therefore, the rest of this discussion is merely speculative rather than conclusive, although it can provide additional information to support previously reported evidence.

First, the absence of an effect of vmPFC stimulation could suggest that other regions may influence the selection component of motor control. As mentioned previously, early sensory areas such as visual cortices are possible candidates (Anderson, 2016; Goard and Dan, 2009; Pinto et al., 2013; Tosoni et al., 2013). However, prefrontal regions show a very complex hierarchical organisation for reward information processing (Hunt and Hayden, 2017), and other possibilities should not be overlooked. It could be for instance that other well-known reward-processing centres located in the prefrontal areas are involved in processing the selection aspects of motor control, such as the cingulate cortex (Blair et al., 2013; Klein-Flugge et al., 2016; Tosoni et al., 2013), which is unfortunately not a possible target for cTBS stimulation due to its deep anatomical location. Another possibility is that vmPFC is indeed involved in the selection process, but that the processing network allows for some compensatory activity, meaning that perturbing vmPFC activity does not affect the network capacity as a whole. Finally, it could be that vmPFC is involved in selection but cTBS is not as effective in perturbing neural activity in vmPFC as in other regions. To our knowledge, only one study reports a significant effect of repetitive cTBS on vmPFC (Lev-Ran et al., 2012), suggesting that perturbation of neural activity with this technique remains possible—though it cannot be ascertained whether our specific stimulation protocol or task design can do so successfully. While that study stimulated participants every 15 minutes, the experiment presented here lasted about 15 minutes as well, suggesting that an effect would have sustained for sufficient time after stimulation ceased. Overall, it is not clear based on our results whether the reliably observed inhibitory effects triggered by M1 cTBS (Huang et al., 2005) can generalise to vmPFC stimulation.

The situation is less ambiguous regarding the absence of an effect of cTBS stimulation on SMA. First, there are numerous studies showing cTBS influences SMA activity (Arai et al., 2011, 2012; Matsunaga et al., 2005; Shirota et al., 2012; Zenon et al., 2015), some of them showing that stimulation can also modulate downstream regions such as M1 (Arai et al., 2011, 2012; Matsunaga et al., 2005; Shirota et al., 2012). This last point indicates that any cTBS effect should be strong enough to lead to consequences even in regions that were not directly stimulated. Additionally, the non-conclusive trend we observe in the peak velocity posteriors with SMA stimulation (figure 3A), and the altered speed-accuracy function (figure 4E) are both in line with the possibility of a global cTBS effect on action vigour – though a larger sample size may be required to reliably expose it. However, due to the “drawer effect” bias (Open Science Collaboration, 2015), it is difficult to ascertain to which extent cTBS stimulation can reproducibly perturb neural processing of SMA. Nevertheless, considering the large set of available studies showing a significant effect of cTBS, and the inconclusive results we report of cTBS on peak velocity and speed-accuracy functions, it is more plausible that other regions implement reward-driven effects on execution, rather than to assume that cTBS is ineffective in manipulating SMA activity. Mainly, the pre-motor area and M1 represent potential alternative candidates. The premotor area is central to movement planning and several studies have shown its sensitivity to reward (Ramkumar et al., 2016; Roesch and Olson, 2003, 2004). Regarding M1, a large literature demonstrates effects of reward on various aspects of M1 processing (Bundt et al., 2016; Galaro et al., 2019; Kapogiannis et al., 2008; Mawase et al., 2016, 2017; Ramkumar et al., 2016; Thabit et al., 2011), making it a suitable candidate for mediating the reward-driven effects observed in our study. Furthermore, we show in Codol et al. (2019) that some execution improvements may be due to an increase in feedback control, likely transcortical (Omrani et al., 2016; Pruszynski et al., 2011) and visuomotor feedback (Carroll et al., 2019). Interestingly, transcortical feedback relies on M1 modulation (Pruszynski et al., 2011), in line with the possibility that M1 supports reward-driven improvements in execution.

Overall, this study shows that the reward-driven effects on reaching are robust and replicable across multiple sessions for a given participant. However, cTBS on the vmPFC and SMA was ineffective in manipulating these effects. While it is difficult to interpret this absence of cTBS effects, we outline possible explanations for this. Notably, the absence of effect following SMA cTBS further bolsters the possibility that reward impacts motor execution at a late stage of the sensorimotor loop, likely at the level of the premotor area or M1.

## Author Contributions

OC, JMG and PJH conceived and designed the research, OC and RJ acquired the data, RJ analysed the imaging data, OC analysed the behavioural data, OC, JMG and PJH interpreted the results, OC wrote the first draft, OC, JMG and PJH edited the manuscript, OC, JMG, RJ and PJH approved the final version of the manuscript.

## Acknowledgments

This work was supported by the European Research Council grant MotMotLearn 637488.

